# Micro-encapsulation extends mycelial viability of *Streptomyces lividans* 66 and increases enzyme production

**DOI:** 10.1101/204677

**Authors:** Boris Zacchetti, Agathoklis Andrianos, Dino van Dissel, Evelien de Ruiter, Gilles P. van Wezel, Dennis Claessen

**Author notes:** These authors contributed equally to this work (Boris Zacchetti, Agathoklis Andrianos, Dino van Dissel). **Contacts:** Boris Zacchetti Agathoklis Andrianos Dino van Dissel Evelien de Ruiter Gilles P. van Wezel Dennis Claessen.

## Abstract

**Background:** Filamentous bacteria of the genus *Streptomyces* produce a large arsenal of industrially relevant antibiotics and enzymes. The industrial production of these molecules occurs in large fermenters, where many streptomycetes form dense mycelial networks called pellets. Pellets are characterized by slow growth and inefficient nutrient transfer and therefore regarded as undesirable from the perspective of productivity. Although non-pelleting strains have increased growth rates, their morphology also leads to a dramatic increase in the viscosity of the culture broth, which negatively impacts the process dynamics.

**Results:** Here, we applied immobilization of *Streptomyces lividans* 66 using alginate as semi-solid matrix. This alginate-mediated micro-encapsulation increased the production of the extracellular enzyme tyrosinase more than three-fold. The increased production was accompanied by extended viability of the mycelium and a dramatic reduction in the release of intracellular proteins into the culture broth.

**Conclusions:** Our data demonstrate the utility of microencapsulation as a powerful technique to achieve higher yields and lower downstream-processing costs of streptomycetes.

## Background

Filamentous organisms are widely used in the field of industrial biotechnology. Of particular relevance are the streptomycetes, multicellular bacteria that produce a vast array of useful metabolites, including over half of the clinically relevant antibiotics, various antitumor agents, immunosuppressants, anthelminthics, antifungals, herbicides and insecticides [1, 2]. In addition, streptomycetes produce and secrete a wealth of extracellular hydrolases, which they employ to degrade the big majority of natural occurring polymers [3]. Streptomycetes grow as filaments (hyphae) that occasionally branch and establish extended cellular networks called mycelia. Growth under industrial settings is characterized by the formation of dense mycelial networks called pellets [4, 5], a phenomenon posing significant drawbacks in terms of industrial applicability. More specifically, pellets only actively grow at the periphery, which limits productivity [6, 7]. Recent work has indicated that it is possible to circumvent pellet formation in *Streptomyces lividans* by interfering with the biosynthesis of extracellular glycans produced by the *cslA-glxA* and *matAB* loci [6, 8]. These glycans mediate the adherence of hyphae, hence leading to the formation of dense clumps and pellets [4]. Deletion mutants of these genes do not form pellets and grow in a dispersed manner. This increases growth and enzyme production rates [6], but comes with the offset of a higher viscosity of the culture broth (our unpublished data). To further complicate matters, pelleted growth appears to be essential at least for the production of some antibiotics [9–11]. All in all, the mycelial mode-of-growth of streptomycetes results in production processes that are characterized by a complex rheology. This translates into suboptimal mass-transfer properties, heat transfer problems, mechanical and oxidative stress [5, 10, 12].

An attractive alternative to classical fermentations is offered by micro-encapsulation, via the immobilization of cells in a semi-solid scaffold, often sodium alginate [13]. The behavior of a number of micro-organisms has been characterized in this immobilized state, which was found to bear several advantages. In comparison to free-living cells, immobilized cells are better protected from harsh environmental conditions [14–16] and enhanced productivity has been reported [17] [18]. Additionally, immobilized cells are readily recycled or filtered, which reduces the yield loss associated with the accumulation of biomass and facilitates downstream processing [19]. In this study, we report that micro-encapsulation of the industrial workhorse *Streptomyces lividans* enhances heterologous production and purity of the extracellular enzyme tyrosinase. Our data indicate that microencapsulation provides protection against shear stress, thereby maintaining mycelial viability and integrity. This in turn stimulates production and reduces contaminations with proteins released by cell lysis.

## Results

### Growth of streptomycetes in calcium-alginate microcapsules extends the viability of the mycelium

To study the effect of microencapsulation on the growth of streptomycetes, spores of *S. lividans, S. coelicolor, S. venezuelae* and *S. griseus* were immobilized in alginate microcapsules (see Materials and Methods). Following the immobilization step, the encapsulated spores were grown in liquid YEME or NMMP_mod_. After 48 h, abundant mycelial growth was detected for all strains in both media (Fig. 1 and Fig. S1). In YEME medium, the encapsulated mycelium of all strains formed highly compact mycelial clumps, while portions of the mycelium that had started to grow out of the microcapsules adopted a more relaxed morphology, whereby individual hyphae could be discerned (Fig. S1). In NMMP_mod_. medium, the encapsulated mycelium formed less compact clumps (Fig. 1). The mycelium of all strains started to grow outside of the microcapsules after 48 h of growth, and became visible as ‘spikes’ protruding from their edges. With the exception of *S. coelicolor,* non-encapsulated mycelium was found in all strains in the liquid during the late stages of growth (Fig. 1 and Fig S2). This phenomenon was particularly evident in *S. griseus* and *S. venezuelae* at 48 h of growth, while it became apparent in *S. lividans* at 96 h of growth (Fig. S2).

**Figure 1.**
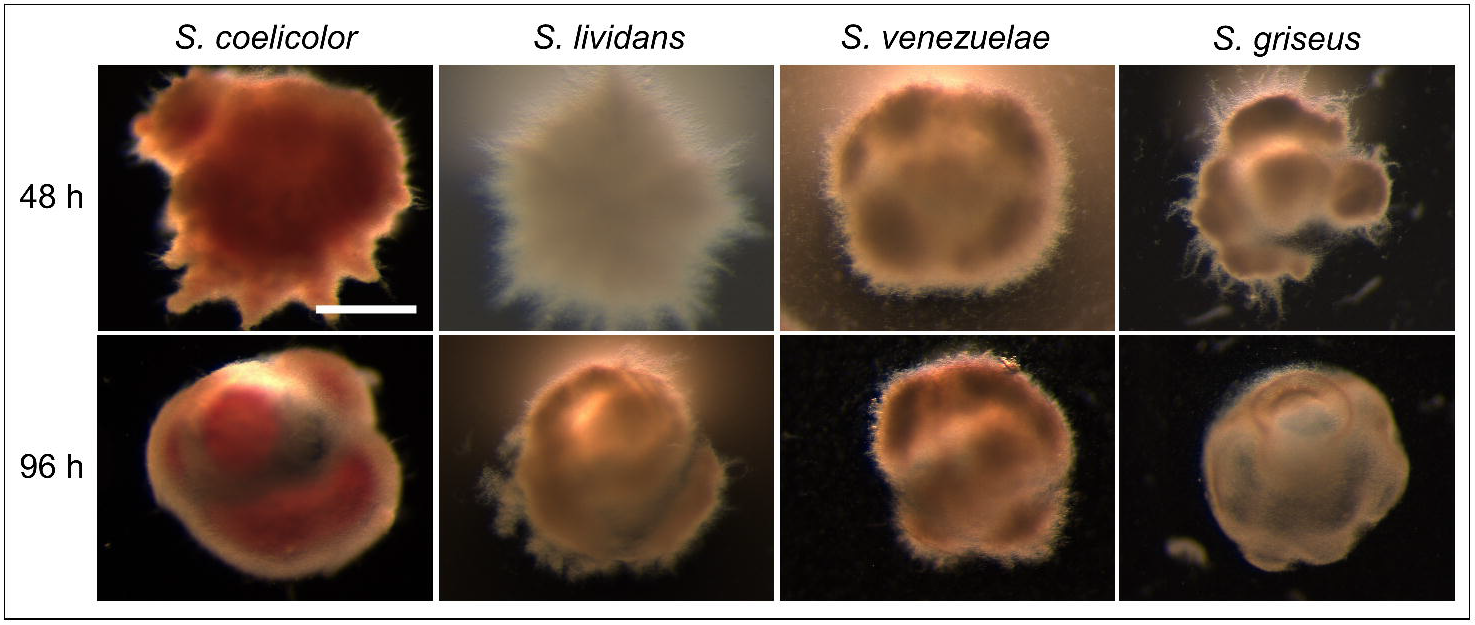
Morphology of encapsulated streptomycetes in NMMP_mod_ medium. Microscopy images of microcapsules of *Streptomyces coelicolor*, *Streptomyces lividans*, *Streptomyces venezuelae* and *Streptomyces griseus* grown in NMMP_mod_ medium at 48 (top panel) and 96 h (lower panel). The scale bar corresponds to 200 μM.

The relaxed morphology of the encapsulated mycelium in NMMP_mod_ compared to YEME medium, which we anticipated to be beneficial for enzyme production, was a reason to further only use NMMP_mod_ medium. We also focused on *S. lividans*, given the prominent role of this strain for the industrial production of enzymes. To analyze how encapsulation affects mycelial viability, we performed a LIVE/DEAD analysis on the encapsulated and free-living mycelium (Fig. 2). Interestingly, a major fraction of the mycelium present in and associated with the microcapsules after 48 and 72 h of growth was viable, as derived from the pronounced staining with Syto-9. In contrast, the presence of damaged mycelium, indicated by the red PI stain, occurred significantly earlier in the free-growing mycelium, either cultivated in the presence or absence of a metal coil (used to counteract aggregation and induce fragmentation by shear). When free-growing mycelium of *S. lividans* was cultivated in the absence of a metal coil in the culture flask, we noticed that the occurrence of damaged mycelium was evident after 48 h of growth, and apparently delayed in comparison to the mycelium suffering from coil-induced shear.

**Figure 2.**
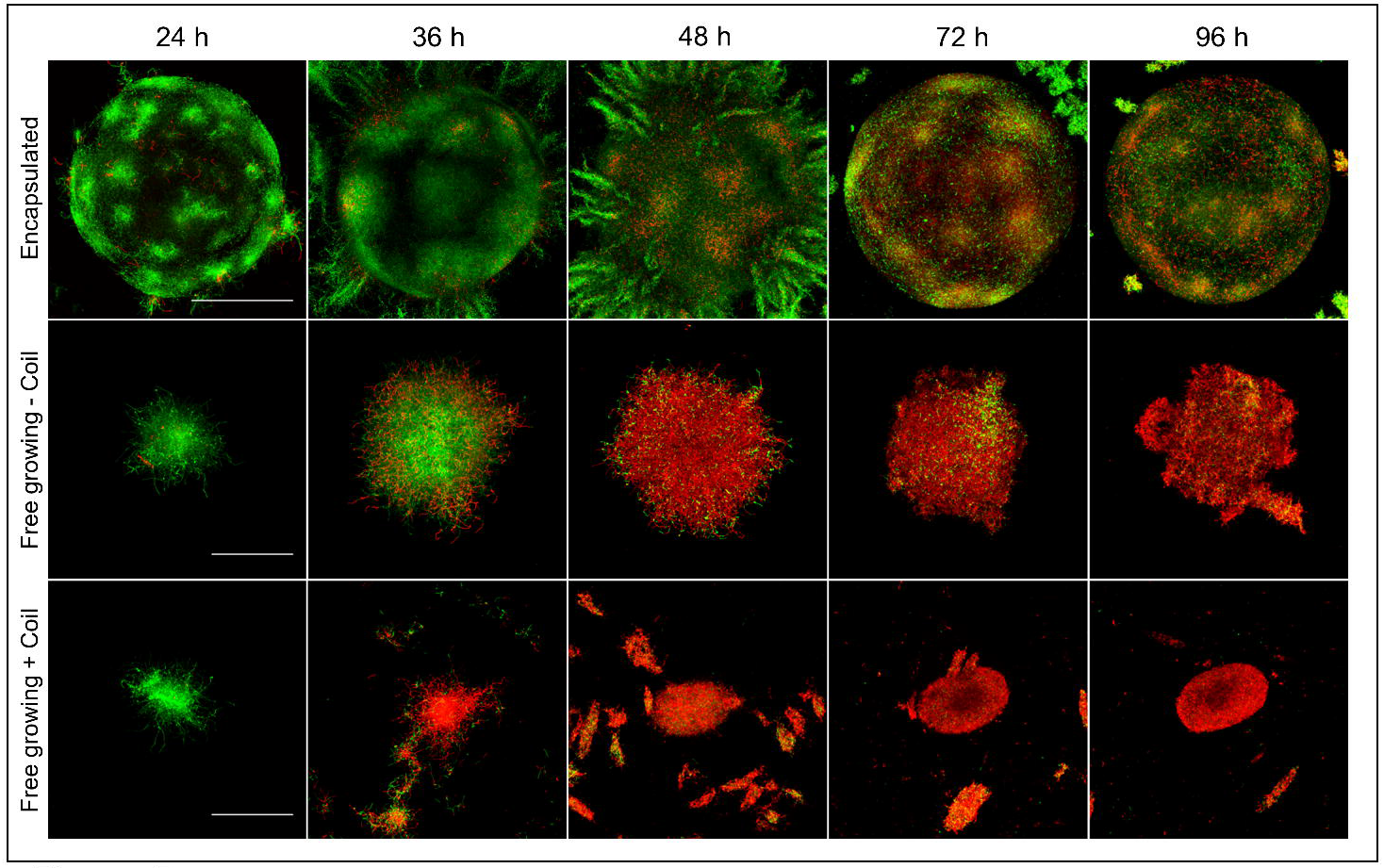
Microencapsulation reduces mycelial damage. LIVE/DEAD staining of *S. lividans* grown in microcapsules (top row), or non-encapsulated in the absence (middle row) or presence (bottom row) of a metal coil. Mycelium stained with Syto9 (green) represents viable mycelium, while propidium iodide-stained mycelium (red) is damaged. Whereas abundant viable mycelium is visible in the encapsulated state at 48 h of growth, the non-encapsulated mycelium appears highly damaged. Note that the mycelium grown in the presence of the coil appears already damaged after 36 h. The scale bar represents 200 μM (top panel) or 100 μM (middle and bottom panel).

### Microencapsulation increases the heterologous production of tyrosinase

To test the effect of microencapsulation on heterologous enzyme production, we introduced plasmid pIJ703 into *S. lividans* 66 [20]. This plasmid contains the *melC2* gene of *Streptomyces antibioticus*, encoding an extracellular tyrosinase that is secreted via the twin arginine translocation pathway [21]. Transformants were selected based on their ability to form the pigmented compound melanin; one of these, hereinafter called *S. lividans* pIJ703, was selected for further analysis.

S. *lividans* pIJ703 was encapsulated, after which the tyrosinase production was assayed and compared to the non-encapsulated controls (Fig. 3). Significantly enhanced activity was detected in the supernatant when *S. lividans* pIJ703 was grown in microcapsules, with a more than three-fold increase in comparison to the non-encapsulated strain. The highest tyrosinase activity in the supernatants of the encapsulated strain peaked after 48 h of growth, followed by a slow and gradual decrease. However, significant tyrosinase activity was still detectable at 72 h of growth. In the non-encapsulated state, the tyrosinase activity peaked at approximately 34 h of growth, after which a rapid decline was detected. After 50 h, tyrosinase activity was barely detectable. Given that the growth rate of the encapsulated mycelium could not be assessed, we measured glucose consumption over time (Fig. S3). The consumption of glucose did not show significant differences between the three culture types, suggesting that the mycelia grew at comparable rates.

**Figure 3.**
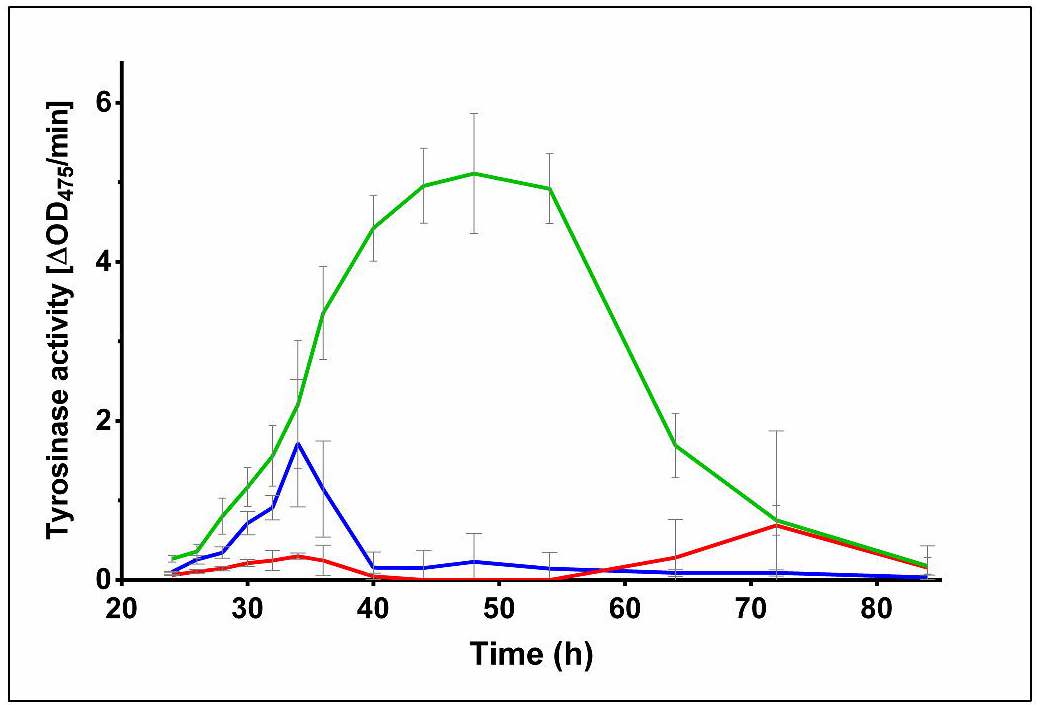
Microencapsulation increases tyrosinase activity in the supernatant. Lines represent the tyrosinase activity present in the supernatant of *S. lividans* pIJ703 grown encapsulated (green), or non-encapsulated in the absence (red) or presence (blue) of a coil. The highest activity was observed in the culture broth of the encapsulated mycelium after 48 h of growth.

We also qualitatively analyzed the extracellular proteins present in the culture supernatants (Fig. 4A). An SDS-PAGE analysis indicated that the supernatant of the encapsulated strain contained an abundant protein with an apparent molecular weight equal to that of tyrosinase (~ 30 kDa) after 48 h of growth, corresponding to the time point where most tyrosinase activity was detected. Conversely, the protein profiles in the supernatants of the nonencapsulated control cultures were more complex and showed a large number of proteins (Fig. 4A, Fig. S4). Western analysis was used to verify that the dominant protein in the supernatant of the encapsulated strain was in fact tyrosinase (Fig. 4B). Consistent with the measured activities in the supernatant, only small amounts of tyrosinase were detected in the non-encapsulated strains. Taken together, these data demonstrate that encapsulation enhances the production of heterologously expressed tyrosinase.

**Figure 4.**
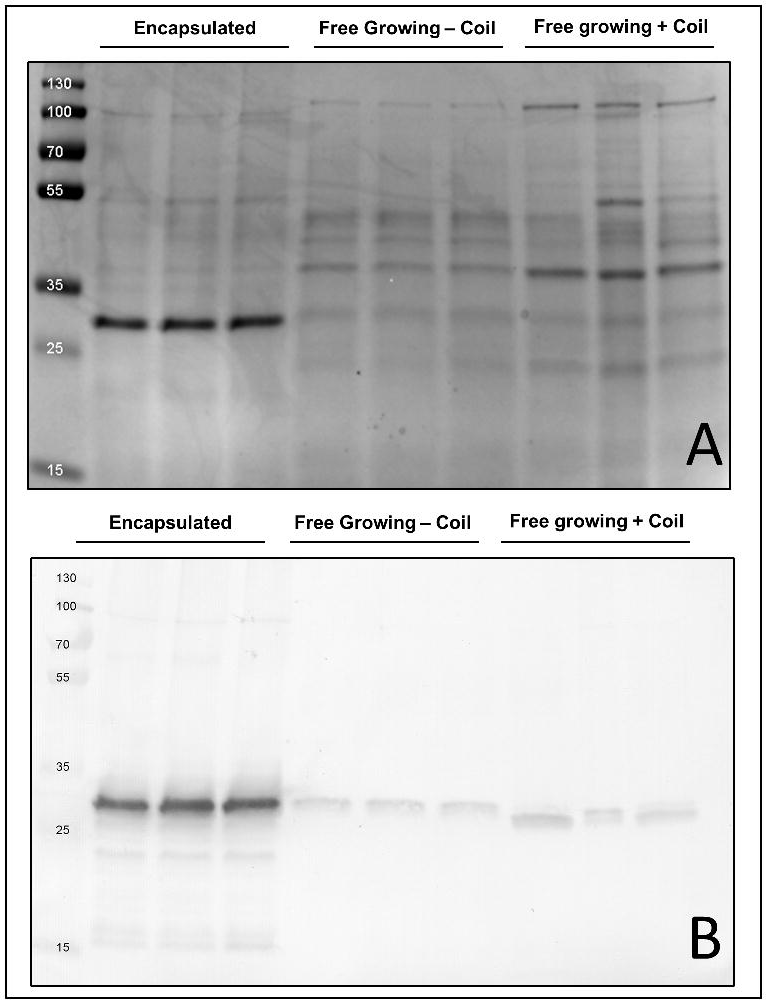
Microencapsulation increases tyrosinase purity in the culture broth. **A)** SDS-page showing the protein profiles in the supernatants of cultures of *S. lividans pIJ703*, grown encapsulated (lanes 2-4), or non-encapsulated in the absence (lanes 5-7) and presence (lanes 8-10) of a metal coil. All cultures were performed in triplicate. Molecular weight markers (lane 1) are indicated in kDa. **B)** Western analysis of the abovementioned supernatants using an anti-Tyrosinase antibody. Molecular weight markers are indicated in kDa.

## Discussion

Streptomycetes are proficient producers of enzymes and antibiotics. For industrial production processes, these organisms are usually grown as liquid-grown cultures in large scale fermenters. The growth of streptomycetes under these conditions is marked by the formation of mycelial particles that consist of interconnected hyphae [4, 22]. Industrial fermenters are typically stirred at high speeds to provide homogeneous mixing but also to ensure that sufficient oxygen and nutrients are available to the growing biomass. This vigorous mixing comes at the cost of severe shear stress, which can cause fragmentation and lysis of the mycelium [23–25]. The concomitant release of intracellular contents into the culture broth thereby complicates product purification [26].

We here present the utility of microencapsulation as a valuable alternative approach circumventing some of the negative aspects of classical fermentations. Microencapsulation physically separates a large fraction of the mycelium from the liquid environment, with the exception of the small mycelial fragments that grow out of the capsules at late time points. While the calcium-alginate is permeable to small molecules [27], the encapsulated mycelium is protected from extrinsic mechanical stress. Our experiments demonstrate that the viability of the mycelium is prolonged inside the microcapsules, which is in agreement with earlier observations in *S. coelicolor [18].* We conclude that this effect is obtained by reducing the degree of shear stress encountered by the mycelium. The earlier occurrence of dead mycelium in non-encapsulated cultures performed with metal coils as compared to those without is also indicative of this fact. Besides extending viability, microencapsulation dramatically increased the production and purity of heterologously produced tyrosinase. The higher production was not only evident from the amount of this protein in the supernatant, but also from measurement of its specific activity throughout growth. More specifically, the amount of active tyrosinase was more than three-fold increased when the mycelium was grown inside the microcapsules.

Shear-induced cell lysis can be a major cause for the dissipation of substrate in streptomycetes [24]. This, accompanied by the observation that the trend in carbon consumption was similar under all conditions, suggests that the encapsulated mycelium invests more energy in production rather than in other processes, such as those related to cell repair and maintenance. The higher purity of the extracellular tyrosinase not only supports that micro-encapsulation reduces cell lysis, but also poses another major benefit: the decreased number of contaminants facilitates product purification and reduces downstream processing costs. Reduced lysis may also prevent the release of intracellular proteases in the culture broth, some of which may lead to the degradation of the desired product. Although we did not analyze this in detail, this aspect may have also contributed to the overall increase of active tyrosinase in the culture broth of the encapsulated *S. lividans* pIJ703. Also, this phenomenon might also explain the decrease in tyrosinase activity in all cultures concomitant with the appearance of abundant dead mycelium.

## Conclusions

Our work demonstrates that micro-encapsulation of streptomycetes extends mycelial viability and enhances the production and purity of enzymes. One issue to overcome is the need to scale up to allow larger scale production with encapsulated strains. Considering this, we anticipate that our approach might be particularly suitable for the production of high-value natural products and enzymes by streptomycetes and possibly other filamentous organisms.

## Methods

### Strains and culture conditions

*Streptomyces coelicolor* A3(2) M145 [28], *Streptomyces lividans* 66 [20] and *Streptomyces venezuelae* diversa were obtained from the John Innes Centre strain collection, and *Streptomyces griseus* DSM40236 from the Deutsche Sammlung von Mikroorganismen und Zellkulturen (DSMZ). MS agar plates [29] were used to prepare spore suspensions of *Streptomyces* strains and to determine colony forming units (CFU) for the spore stocks. For liquid-grown cultures, YEME medium [29] or a modified NMMP medium (NMMP_mod_) were used. The buffer system of NMMP_mod_ was optimized to avoid the detrimental effect of phosphates on the integrity of the alginate microcapsules [30]. For the preparation of 1 liter NMMP_mod_ medium, 100 ml 0.25 M TES buffer (pH 7.2), 10 ml 0.1M Na-K buffer (pH 6.8), 25 ml 20% glucose and 65 ml milliQ water were added to 800 ml NMMP basis [29]. For experiments using strains containing the tyrosinase-expressing plasmid pIJ703 [21], 10 μM CuSO_4_ were added to the growth medium. All cultures were grown in a total volume of 100 ml of liquid medium contained in 250 ml Erlenmeyer flasks. Cultures were grown in an orbital shaker set at 30° C and 160 rpm. Unless differently stated, all experiments were performed in duplicate. For micro-encapsulation experiments, on average 75 viable spores were incorporated into every capsule with an average diameter of 415 μm. To this end, spores were suspended in sterile liquid sodium alginate and thoroughly mixed before the preparation of microcapsules. A total of 5 ml of alginate microcapsules was used to inoculate 100 ml of medium (*see below*). We calculated the equivalent number of spores, which we used to inoculate all other cultures and which corresponded to 10^5^ spores per ml of medium.

### Microscopy

A Leica MZ12 stereo microscope was used for the visualization of microcapsules and encapsulated mycelium. For the visualization of live and dead mycelium, samples were stained with Syto-9 and propidium iodide (PI) (Invitrogen). To this end, pellets and microcapsules were briefly sedimented via centrifugation (10 min at 2000 rpm at room temperature) and resuspended in PBS, to which Syto-9 and PI were added to a final concentration of 5 μM and 15 μM respectively. After mixing and incubating for 10 minutes in the dark at 30° C, samples were analyzed using a Zeiss LSM 5 EXCITER confocal microscope. Stained samples were excited at 488 and 543 nm for Syto-9 and PI, respectively. The fluorescence emission of Syto-9 was monitored in the region between 505-545 nm, while a long-pass filter of 560 nm was used to detect PI [31]. The pictures shown in Fig. 2 represent Z-stacks of at least 15 layers in the specimen with a slice thickness of 7 μm for microcapsules and 4 μm for pellets.

### Encapsulation of *Streptomyces* spores in calcium alginate

Sodium alginate (Sigma-Aldrich, CAS:9005-38-3) was dissolved under constant stirring for 1 hour in milliQ water to obtain a 2% solution. To remove undissolved micro-particles and other contaminants, the obtained solution was passed through two different filters. The first filter had a pore size of 1.2 μm (GE Healthcare, CatNo:1822-047) and was used in a vacuum filtration apparatus (PALL Magnetic Filter Funnel). The filtrate was then filter-sterilized using a syringe filter with a pore size of 0.22 μm (Sarstedt). For the production of calcium-alginate microcapsules, a home-made device was used similar to that described in **[32]**. This apparatus is based on a coaxial gas-flow extrusion principle, with sterile air as the used gas. The air flow was regulated via a controller (Kytola, Model E) and was set at 3 liters per minute, thus yielding alginate particles with an average diameter of 415 μm (± 12.3 μm; based on analyzing 150 particles). A constant alginate flow was obtained by using a syringe pump (Fisher-Scientific) set at 30 ml/hour. The microcapsules were produced by dispersing the extruded alginate using a home-made nozzle that allowed co-axial laminar flow. While the alginate constitutes the inner sheath, air is flown in a co-axial fashion and determines the rate of formation of the alginate drops. This allowed control over the volume of the falling droplets, which were collected into a gently stirred 200 mM CaCl_2_ solution. The cross-links formed through sodium/calcium ion exchange almost instantly transformed the liquid drops into gel-like microcapsules. The alginate microcapsules were left to harden in the stirring 200 mM CaCl_2_ solution for 5 minutes. The obtained suspension of calcium-alginate microcapsules was then filtered using a vacuum filtration apparatus (PALL, GE Healthcare filters as above), after which the microcapsules were washed three times with 500 ml sterile demi-water.

### Glucose measurement assay

Glucose concentrations were determined using a commercial kit (Megazyme, HK/G6P-DH), according to the instructions of the manufacturer.

### Tyrosinase activity assay

The specific activity of tyrosinase produced by *S. lividans* harboring pIJ703 was determined by following the conversion of L-3,4-dihydroxyphenylalanine spectrophotometrically at a wavelength of 475 nm, as described earlier [33].

### SDS-PAGE and Western blot analyses

Supernatants of liquid-grown cultures were harvested after 48 h of growth. The culture samples were first centrifuged for 10 min at 5000 rpm and 4° C, after which the supernatants were filtered through 0.22 μM syringe filters (Sarstedt), to remove any possible contaminants (e.g whole cells, spores). Extracellular proteins were concentrated via acetone precipitation. Briefly, 1.2 ml of cold acetone (-20°C) were added to 300 μl of supernatant sample. Following thorough mixing, the sample was kept at -20°C for 1 hour and subsequently centrifuged at 13,000 rpm for 10 min at 0°C. Subsequently, the liquid was removed without disturbing the protein pellet, after which 500 μl of cold acetone were added. After a second centrifugation step, the acetone was removed and the pellet was dried at 37 °C for 10 min. The obtained protein pellets were dissolved in 30 μl of 10 mM Tris-HCl buffer (pH 8.0). A Bradford analysis was used to determine the protein concentrations in the obtained samples, and 2 μg of proteins were used for separation by SDS-PAGE on precast 12 % miniprotean TGX Gels (BioRad) at 205 V, 200 mA for approximately 50 min. Proteins were transferred to polyvinylidene difluoride (PVDF) membranes (GE Healthcare) and incubated overnight with anti-Tyrosinase polyclonal antibodies (1:25,000 dilution). Following 1 hour of incubation with goat anti-rabbit alkaline phosphatase, detection was carried out with NBT/BCIP. The relative quantification of proteins on SDS-pages was performed using ImageJ (version 1.48f).

## Declarations

**Ethics approval and consent to participate:** Not applicable

**Consent for publication:** Not applicable

**Availability of data and material:** Not applicable

**Competing interests:** The authors declare no competing interests

**Funding:** This work was supported by grant 12957 from the Dutch Applied Research Council to DC

**Authors' contributions:** BZ performed the experiments with the help of AA, DvD and ER. DC, BZ and DVD conceived the study. BZ and DC wrote the manuscript with the help of GPvW. All authors read and approved the final manuscript.

## Acknowledgements

Not applicable

## Figure Legends

**Figure S1: Morphology of encapsulated streptomycetes in YEME medium.** Microscopy images of microcapsules of *Streptomyces coelicolor, Streptomyces lividans, Streptomyces venezuelae* and *Streptomyces griseus* grown in YEME medium for 48 (top panel) and 96 h (lower panel). No scale bar is added since not all pictures are taken using the same magnification (mainly to allow the visualization of the protruding mycelium). As a reference, the microcapsules have an average size of 415 μm.

**Figure S2: Growth and detachment of mycelium from microcapsules containing different streptomycetes.** Overview images of the mycelium of *Streptomyces lividans, Streptomyces griseus* and *Streptomyces venezuelae* grown in NMMP_mod_ medium for 48 (top panel) and 96 h (lower panel). Note that detached mycelial fragments are evident in the culture broth of *S. griseus* and *S. venezuelae* at 48 h. After 96 h, detached mycelial fragments are also observed in *S. lividans*. The scale bar corresponds to 500 μm.

**Figure S3. Glucose consumption by encapsulated and non-encapsulated mycelium.** The residual glucose concentrations (in g/L) in NMMP_mod_ medium are shown when *Streptomyces lividans pIJ703* is grown in micro-capsules (green), or non-encapsulated in the absence (red) and presence (blue) of a metal coil.

